# Secretory and transcriptomic responses of mantle cells to low pH in the Pacific oyster (*Crassostrea gigas*)

**DOI:** 10.1101/2023.01.19.524809

**Authors:** Nicolás Zúñiga-Soto, Ingrid Pinto-Borguero, Claudio Quevedo, Felipe Aguilera

## Abstract

Since the Industrial Revolution, the concentration of atmospheric carbon dioxide (CO_2_) due to anthropogenic activities has increased at unprecedented rates. One-third of the atmospheric anthropogenic CO_2_ emissions are dissolved in the oceans affecting the chemical equilibrium of seawater, which in turn leads to a decrease in pH and carbonate ion (CO_3_^2−^) concentration, a phenomenon known as ocean acidification (OA). This chemical disequilibrium can be detrimental to marine organisms (e.g., mollusks) that fabricate mineralized structures based on calcium carbonate (CaCO_3_). Most studies on the effect of reduced pH in seawater have been conducted on the early developmental stages of shell-building invertebrates, neglecting how adult individuals face OA stress. Here, we evaluate histological, secretory, and transcriptional changes in the mantle of adult oysters (*Crassostrea gigas*) exposure to ambient (8.0 ± 0.2) and reduced (7.6 ± 0.2) pH during 20 days. Most histological observations did not show differences in terms of mantle cell morphology. However, Alcian Blue/PAS staining revealed significant differences in the number of Alcian Blue positive cells in the mantle edge, suggesting a decrease in the secretory activity in this morphogenetic zone. Transcriptomic analysis revealed 172 differentially expressed genes (DEGs) between mantle tissues from adult oysters kept in normal and reduced pH conditions. Almost 18% of the DEGs encode secreted proteins that are likely to be contributing to shell fabrication and patterning. 17 of 31 DEGs encoding secreted proteins correspond to oyster-specific genes, highlighting the fact that molluscan shell formation is underpinned by a rapidly evolving secretome. The GO analysis of DEGs encoding secreted proteins showed that they are involved in the cellular response to stimulus, response to stress, protein binding, and ion binding, suggesting these biological processes and molecular functions are altered by OA. This study demonstrates that histology and gene expression profiling can advance our understanding of the cellular and molecular mechanisms underlying adult oyster tolerance to low pH conditions.

## 1 Introduction

Since the beginning of the Industrial Revolution, carbon dioxide (CO_2_) levels in the atmosphere have been increasing from 280- to 400-ppmv due to human activities (Falkowski et al., 2000). Most global CO_2_ emitted from anthropogenic activities is eventually absorbed by the oceans, producing a progressive increase in ocean inorganic carbon concentrations, which in turn leads to changes in the carbonate chemistry through a process called ocean acidification (OA) (Caldeira and Wickett, 2003;Doney et al., 2009). Consequently, the acid-base chemistry of seawater tends towards more acidic conditions, reducing the pH and saturation state of carbonate minerals (Doney et al., 2009;Hurd et al., 2019). The subsequent increase in carbonic acid reduces the availability of carbonate ions (CO_3_^2−^) in oceans, thus affecting marine organisms that use calcium carbonate (CaCO_3_) as building blocks to fabricate their shells (Orr et al., 2005;Fabry et al., 2008;Guinotte and Fabry, 2008;Gazeau et al., 2013;Hurd et al., 2019). Ocean acidification has been reported to affect ocean biota, but the effects of reduced pH depend greatly on the marine organisms studied (Ries et al., 2009). For instance, some marine organisms are resilient and can acclimate to OA (Kroeker et al., 2010;Munday et al., 2011;McCulloch et al., 2012), while most shell-building invertebrates, including corals, echinoderms, and mollusk face trouble fabricating their shells in the presence of elevated CO_2_ levels and reduced pH in seawater (Gazeau et al., 2007;Kurihara et al., 2007;Miller et al., 2009;Talmage and Gobler, 2009;Todgham and Hofmann, 2009;Albright et al., 2010;Brennand Sheppard et al., 2010;Gazeau et al., 2010;Albright and Langdon, 2011;Parker et al., 2011;Gazeau et al., 2013;Parker et al., 2013;Doney et al., 2020). Accumulating evidence suggests that mollusks find it more difficult to deposit their CaCO_3_ shells and suffer a range of negative impacts including smaller-sized embryos and larvae, decreased shell thickness, increased larval developmental time, shell abnormalities, and reduced survival and metamorphosis rates (Beniash et al., 2010;Parker et al., 2013;Ko et al., 2014). This has been used to argue that the early developmental stages of mollusks are more sensitive to ocean acidification than the late (juvenile/adult) stages (Parker et al., 2013). The adult mollusk shell is a remarkably stable organo-mineral biocomposite, in which the mineral phase (CaCO_3_) makes up 95-99% by weight and the remainder is composed of organic macromolecules such as proteins, polysaccharides, and lipids (Marin et al., 2012). Molluscan shells are assembled extracellularly and are the result of the secretory activity of the outer epithelium of the mantle (Kocot et al., 2016;McDougall and Degnan, 2018), which consists of highly specialized cell types (Sud et al., 2002;McDougall et al., 2011;Budd et al., 2014) and is divided into distinct morphogenetic zones (Jolly et al., 2004;Takeuchi and Endo, 2005;Jackson et al., 2006;Jackson et al., 2007); each responsible for the secretion of shell matrix proteins (SMPs) that influences the formation of specific shell layers (Takeuchi and Endo, 2005;Gardner et al., 2011;Marie et al., 2012;Herlitze et al., 2018). Therefore, the control of specific shell layers is provided by the coordinated expression of a suite of genes encoding secreted proteins, the so-called “mantle secretome” (Jackson et al., 2006;Kocot et al., 2016;Aguilera et al., 2017). Previous studies that analyzed the effect of OA on late developmental stages (juvenile/adult) found a reduction in the calcification rate, growth, strength, and increased dissolution of molluscan shells (Parker et al., 2013;Avignon et al., 2020). However, it remains unclear how OA affects the cellular morphology, secretory activity, and gene expression profiles of the outer epithelium of the mantle tissue in adult mollusks.

Given the different physiological, behavioral, and habitat requirements which exist between early (embryonic/larval) and late (juvenile/adult) stages in mollusks (Parker et al., 2013), the effects of OA are likely to differ depending on their stage of development. Therefore, a thorough understanding of the mechanisms that control cellular and molecular response to low pH, at different developmental stages, is needed to uncover the sensitivity of marine organisms exposed to OA. Here, we evaluate histological, secretory, and transcriptional changes in the mantle tissues of adult oysters (*Crassostrea gigas*) exposed to seawater at a pH of 8.0 ± 0.2 and 7.6 ± 0.2 for 20 days to determine the underpinning molecular mechanisms and physiological responses of oysters coping with environmental pH variations, and thus to provide an integrated view of the cellular response and molecular pathways affected by OA.

## 2 Materials and Methods

### Adult oyster maintenance and CO_2_ perturbation setup

Adult oysters (*Crassostrea gigas*) were purchased from a culture farm located in Coliumo Bay (36.5431°S, 72.9589°W), Concepcion, Chile. Adult oysters were acclimated in 30L tanks with artificial seawater (Instant Ocean Salt) adjusted to 35‰ and constant aeration for two weeks before starting exposure experiments (**Figure 1A**). After acclimation, adult oysters were randomly divided into ten individual 2L flasks with artificial seawater at 35‰ (**Figure 1A**). Two pH treatment levels (pH 7.6 ± 0.2 and pH 8.0 ± 0.2) were targeted with five replicate tanks per treatment (**Figure 1A**). The normal pH seawater was produced by bubbling individual tanks with atmospheric air through air diffusers, whereas low pH seawater was produced by bubbling CO_2_-enriched air until the desired pH value (7.6 ± 0.2). Then, oysters were kept with a low air input to maintain the low pH value. These pH levels were chosen because encompass potential mean oceanic pH levels estimated for current oceans (pH 8.0) and reflect atmospheric CO_2_ for future oceans (pH 7.6) predicted by the Intergovernmental Panel on Climate Change IPCC for the year 2100 (Caldeira and Wickett, 2005). Artificial seawater was renewed every day with pre-bubbled seawater that had been equilibrated previously, according to the experimental trials (pH 8.0 ± 0.2 or pH 7.6 ± 0.2). The pH was measured during each seawater renewal with a pH meter (MW804, Milwaukee). During the acclimation and experimental period, adult oysters were fed daily with a commercial algal mix (PhytoBlast, Continuum), containing a mixture of *Nannochloropsis sp., Tetraselmis sp., Pavlova sp*., and *Thalassiosira sp*. microalgae. After 20 days of cultivation, 10 individuals (6 for histology and secretory assays, and 4 for transcriptomic analysis) were sacrificed. Before the dissection of the mantle tissues, animals were relaxed in 1M MgCl_2_ in filtered artificial seawater.

**Figure 1.**
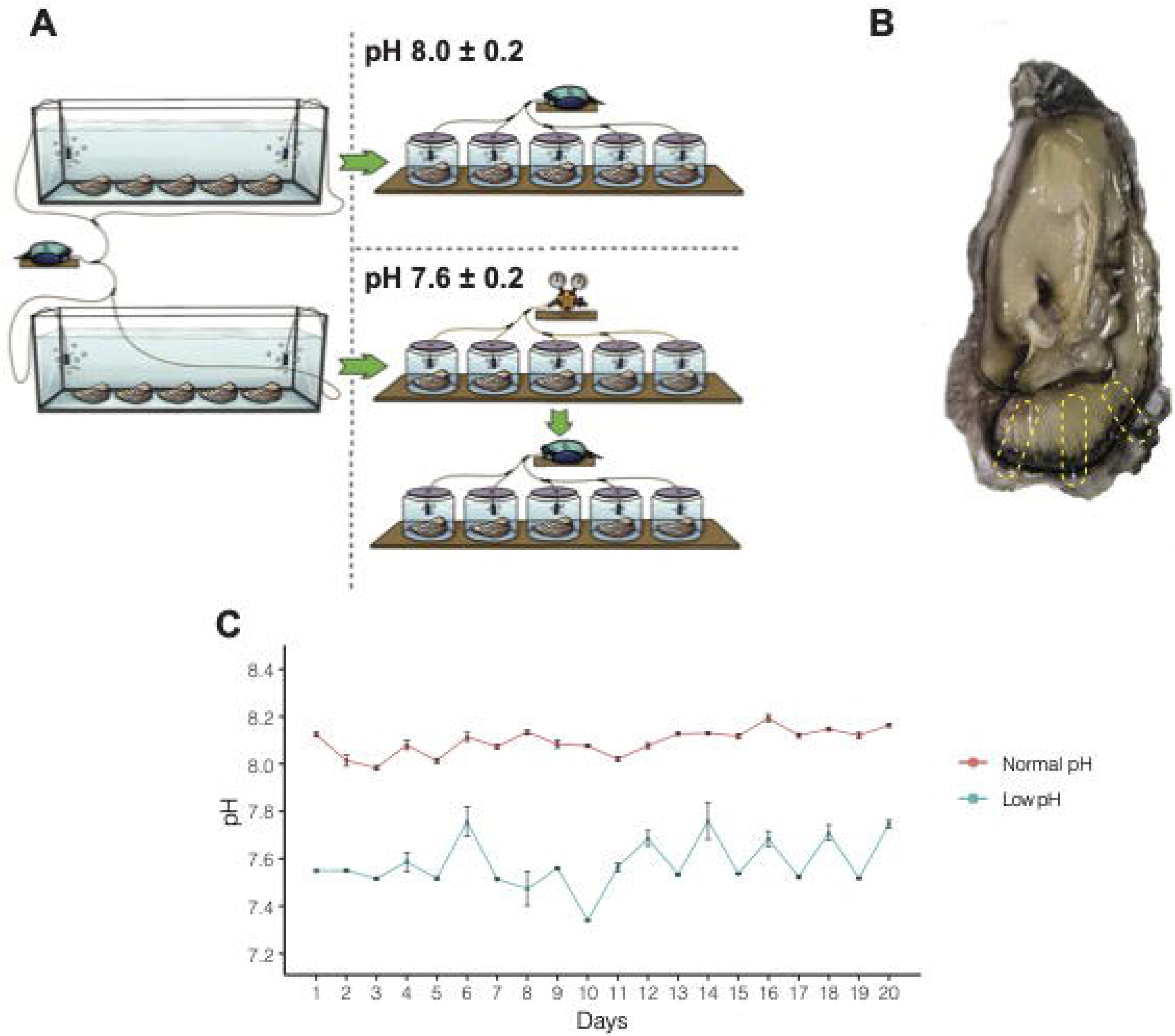
Experimental design and pH measurement to study the effect of OA on adult oysters. **A)** A total of ten oysters were acclimated for two weeks and then split into two experimental treatments. After acclimation, five oysters were kept under normal pH (8.0 ± 0.2) and low pH (7.6 ± 0.2) conditions for 20 days. **B)** After the experimental period, oysters were sacrificed, and different regions of the mantle tissue were removed to conduct histological and transcriptome analyses. **C)** pH on each experimental flask was measured three times daily for 20 days.

### Histological procedures on mantle tissues

Samples from different regions of the mantle tissue (**Figure 1B**) were dissected and fixed in 4% paraformaldehyde in PBS for general histology. Tissue samples were embedded in paraffin and sectioned at 3-5μm using a microtome (Microm HM325, Thermo Scientific). Before histological staining, tissue sections were treated with xylene to remove paraffin, rehydrated through a graded ethanol series, and running tap water. To characterize mantle histology, tissue sections were stained using the hematoxylin and eosin (H&E) staining kit (ab245880, abcam) following the manufacturer’s instructions. To characterize the cellular nucleus, tissue sections were stained with toluidine blue as described by Sridharan and Shankar (2012). Briefly, dehydrated tissue sections were treated with 0.1% toluidine blue for 30 minutes, and staining excess was removed through three washes with distilled water. Finally, tissue sections were dehydrated at 60°C to remove distilled water before mounting.

To characterize connective and muscular tissues in the mantle, we used the protocol described by Masson (1929). Briefly, tissue sections were incubated in Bouin solution at 60°C, and Bouin excess was removed using 70% ammoniac-alcohol solution and washed using running tap water. After washing, tissue sections were stained with Weigert’s Hematoxylin for 30 minutes and differentiation was performed using a 70% acid-alcohol solution. Tissue sections were then plunged in Biebrich Scarlet for 3 minutes and treated with phosphotungstic and phosphomolybdic acids for 3 minutes for allowing the binding of aniline blue. Tissue sections stained with aniline blue were differentiated using 1% acetic acid and dehydrated at 60°C before mounting.

After staining procedures, tissue sections were dehydrated using xylene and mounted using DPX mounting media (06522, Sigma). Tissue sections were observed under a microscope Axiolab 5 (Zeiss) and images were processed using ImageJ v1.53 (Schneider et al., 2012) for adjusting contrast and lightness.

### Mantle tissue secretory cells and quantification

To identify possible secretory cells in the mantle tissue, the histochemical procedure based on the periodic acid-Schiff stain (PAS) and Alcian Blue (AB) was used for neutral and acid mucopolysaccharide staining, respectively. We used the AB/PAS staining kit (ab245876, abcam) following the manufacturer’s instructions, with some modifications. Briefly, tissue sections were plunged into 3% acetic acid for 2 minutes; the acetic acid excess was rinsed, and tissue sections were incubated in AB (pH 2.5) for 30 minutes. Staining excess was removed using running tap water and distilled water; then, tissue sections were plunged in the periodic acid solution for 5 minutes, washed in distilled water, and incubated in Schiff’s solution for 20 minutes. Finally, tissue sections were stained using Mayer’s Hematoxylin for 2 minutes, rinsed with running tap water, dehydrated using a graded ethanol series followed by xylene, and mounted with DPX mounting media (06522, Sigma). Digital images were captured using a microscope Axiolab 5 (Zeiss) equipped with an Axiocam 208 color camera (Zeiss). Images were processed using ImageJ v1.53 (Schneider et al., 2012) for adjusting contrast and lightness.

To calculate the number of secretory cells in mantle tissue exposed to normal and low pH, we performed AB positive cells counting along the outer mantle epithelium as a proxy to infer whether secretory cells might be delivering macromolecules (e.g., SMPs) into the extrapallial space where the interactions between CaCO_3_ and proteins occurs (Huang and Zhang, 2022). The number of secretory cells was counted in the mantle edge and mantle pallial morphogenetic zones of adult oysters kept under normal and low pH conditions. t-test was applied to look for significant differences in the number of secretory cells in either mantle edge or mantle pallial between oysters kept under normal pH (8.0 ± 0.2) or low pH (7.6 ± 0.2). The data were considered statistically significant at p < 0.05. All data were analyzed using R v4.1.3 (R Development Core Team, 2022) and plots were made using ggplot (Wickham, 2016).

### Transcriptomic analysis of mantle tissues exposed to normal and low pH

After relaxation, mantle tissues were dissected and stored immediately at −80°C until further RNA extraction. Total RNA was extracted from mantle tissues of adult oysters kept under normal and low pH conditions with TriReagent (T9424, Sigma) following a modified protocol described by Gao et al. (2001) to remove inhibitory pigments. RNA was quantified using a NanoDrop™ Lite Spectrophotometer (Thermo Scientific) and shipped to Macrogen (Korea) to perform cDNA library preparation and sequencing in an Illumina NovaSeq6000 paired-end mode.

Sequencing quality was performed using FastQC v0.11.9 (Andrews, 2010), and reads were trimmed with Trim Galore! v0.67 (Krueger, 2012). High-quality reads were mapped to the *C. gigas* genome (Zhang et al., 2012) using STAR v2.6.1a_08-27 (Dobin et al., 2013), and those reads that mapped to transcripts were counted with featureCounts v1.6.3 (Liao et al., 2014). Counted reads were also used for identifying differentially expressed genes (DEGs) using edgeR v3.36.0 (Robinson et al., 2010) and an FDR < 0.05 and an absolute value of log2-fold change ≥ 2. DEGs were visualized as volcano plot using the VolcaNoseR web app (Goedhart and Luijsterburg, 2020).

Proteins encoded by those genes identified as differentially expressed were classified as cytosolic, secreted, or transmembrane following the procedure described by Aguilera et al. (2017). Briefly, *C. gigas* proteins were searched for signal peptides using a local installation of SignalP v4.1 (Petersen et al., 2011), with the following parameters: -s best -t euk -u 0.45 -U 0.50. Proteins predicted as non-secretory were classified as cytosolic proteins. Signal peptide-positive proteins were additionally screened for transmembrane domains using a local installation of THMHH v2.0c (Krogh et al., 2001), or for mitochondrial, Golgi, or lysosomal targeting using the TargetP v2.0 web server (Emanuelsson et al., 2007). Positive proteins were classified as transmembrane or organelle targeting, respectively. Transcript counts (normalized using TMM transformation in edgeR) from DEGs encoding secreted proteins were used to generate a heatmap for visualization of differentially expressed genes with the ClustVis web tool (Metsalu and Vilo, 2015). Gene Ontology (GO) annotation of *C. gigas* gene models was performed following the Trinotate pipeline (Haas et al., 2013). Those GO terms associated with DEGs (encoding secreted and non-secreted proteins) were analyzed using WEGO v2.0 (Ye et al., 2018).

DEGs encoding secreted proteins were further investigated whether they (i) were likely to encode shell matrix proteins (SMPs) based upon similarity to proteins that had previously been identified from *C. gigas* shells, or they had transcriptional regulatory or signaling activity roles. Similarity to SMPs was performed by BLASTP searchers (Camacho et al., 2009) against an in-house database of published proteins that had previously been identified from *C. gigas* larval and adult shells (Marie et al., 2011;Zhang et al., 2012;Zhao et al., 2018), using an e-value of 1e^−100^. As many SMPs contain repetitive low-complexity domains (RLCDs) (Jackson et al., 2010;McDougall et al., 2013;Aguilera et al., 2017), BLASTP searches were conducted without filtering for low-complexity regions and without compositional adjustment. Potential transcription factor or signaling activity was ascertained by searching GO term annotations for GO:000370 (DNA-binding transcription factor activity), or the word “signal” in the GO description.

## 3 Results

Ocean acidification is a concerning and global-scale phenomenon with negative impacts on diverse animal groups that form calcified structures based on CaCO_3_, being mollusks one of the groups most affected (Gazeau et al., 2007;Gazeau et al., 2013). Many studies have shown these negative impacts in the early-life stages of mollusk (Parker et al., 2013); however, the impact of ocean acidification on adult individuals has not been well explored yet. To achieve a better understanding of the effects of OA in adult mollusks, we maintained adult oysters under normal (8.0 ± 0.2) and low (7.6 ± 0.2) pH for 20 days (**Figure 1C**). To evaluate the effects of low pH, we employed an integrative approach, combining histological and transcriptomic analyses, to investigate the cellular and molecular responses of adult mollusks exposure to OA condition.

### Ocean acidification does not affect mantle tissue histology

The mantle is composed of epithelia, loose stroma, muscular, adipose, and connective tissues (**Figure 2A**). Histological observations show that epithelial cells are closely aligned to form the epidermis of the mantle that extends into the connective tissue (**Figures 2B and 2C**). H&E and toluidine blue staining reveal numerous invaginations along the outer mantle epithelium covering the morphogenetic zones known as mantle edge and mantle pallial (**Figure 2A and Supplementary Figure 1**). The inner fold is similar in terms of length and shape compared to the other mantle folds (**Figure 2A and Supplementary Figure 1**). The outer fold is composed of a columnar epithelium (**Figures 2B and 2D**), whereas the middle fold is characterized by a pseudostratified columnar epithelium, with nuclei at different heights (**Figure 2D**). This epithelial transition in the periostracal groove may be relevant because this mantle region is responsible for the secretion of the proteinaceous periostracum layer (McDougall et al., 2011;Marin et al., 2012;Kocot et al., 2016).

**Figure 2.**
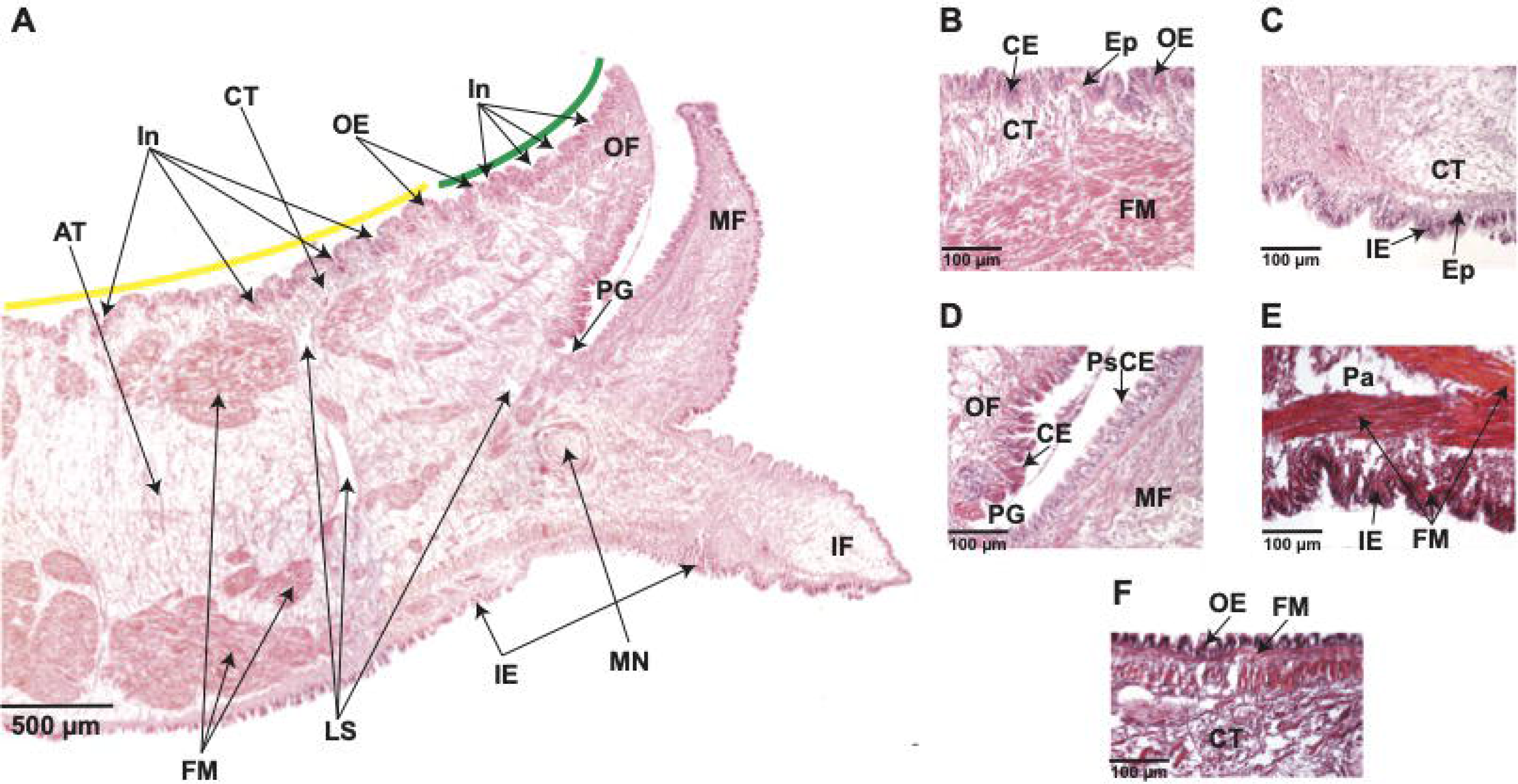
Transverse sections of the mantle tissue of *C. gigas* adult oyster were kept under normal pH (8.0 ± 0.2) conditions. **A)** Hematoxylin and Eosin-stained section through the mantle depicting various structures, including the outer epithelium (OE), outer fold (OF), periostracal groove (PG), middle fold (MF), inner fold (IF), the inner epithelium (IE), mantle nerve (MN), loose stroma (LS), fibers muscle (FM), adipose tissue (AT), and connective tissue (CT). Along the outer epithelium (OE), there are several invaginations (In) covering the mantle zones known as mantle edge (green line) and mantle pallial (yellow line). Scale bar = 500μm. **B)** Hematoxylin and Eosin-stained transverse section through the outer epithelium (OE) showing the presence of aligned columnar epithelial (CE) cells forming the epidermis, which extends into the connective tissue (CT). Scale bar = 100μm. **C)** Inner epithelium (IE) section with Hematoxylin and Eosin staining also shows aligned columnar epithelial (CE) cells that form the epidermis (Ep) extending into the connective tissue (CT). Scale bar = 100μm. **D)** Hematoxylin and Eosin-stained section at the base of the periostracal groove (PE) depicting the presence of columnar epithelium (CE) on the outer fold (OF) side, but not on the middle fold (MF) side that is characterized by a pseudostratified columnar epithelium (PsCE) with nuclei at different heights. Scale bar = 100μm. **E)** Inner epithelium (IE) section stained with Trichome-Masson depicting bundles of fibers muscle (FM) that underlie the inner epithelium (IE) and near the parenchyma (Pa). Scale bar = 100μm. **F)** Trichrome-Masson-stained section showing the presence of fibers muscle (FM) underlying the outer epithelium (OE). Scale bar = 100μm.

The mantle musculature consists of bundles of fibers muscle located in the parenchyma and underneath the outer and inner epithelia (**Figures 2A and 2E-F**). Within the parenchyma, numerous long muscle fibers are densely organized and longitudinally distributed, with apparently no striated (**Figure 2E**). The outer epithelium adjoins a layer of longitudinal muscles, without apparent striated, embedded in connective tissue (**Figure 2F**). These histological observations were similar in the mantle tissues from adult oysters kept in low pH conditions (**Supplementary Figures 2 and 3**), suggesting that ocean acidification does not affect mantle tissue anatomy and cell morphology, at least for 20 days of exposure to reduced (7.6 ± 0.2) pH.

### The distribution of mantle secretory cells is susceptible to ocean acidification

Staining with AB at pH 2.5 show that sulfated mucosubstances (stained in blue) produced by mucus cells are distributed at the base of the epithelium of the three folds of the mantle (**Figures 3A and 3B**). These secretions are equally evident in outer and inner epithelia compared to the epithelium lining the middle fold (**Figures 3A and 3B**). Mucus cells are observed in the mantle edge and mantle pallial zones of adult oysters maintained at normal (8.0 ± 0.2) pH, but with a high concentration of AB-positive (sulfated mucosubstances) cells lining the outer epithelium of the mantle and the near connective tissue (**Figure 3A**). In contrast, there is a low number of mucus cells in the mantle tissues of adult oysters kept at a low (7.6 ± 0.2) pH condition (**Figure 3B**). We did not observe any cells containing AB-positive signal in the connective tissue of adult oysters kept at a reduced (7.6 ± 0.2) pH condition (**Figure 3B**). Remarkably, the difference in terms of the number of secretory (mucus) cells is related to a particular morphogenetic zone of the mantle (n = 54; p 2.2^e-16^) (**Figure 3C**), suggesting a low capacity to secrete macromolecules in the mantle edge region in adult oysters kept under reduced (7.6 ± 0.2) pH condition for 20 days. Given that different secretory repertoire control the formation of prismatic and nacreous layers (Takeuchi and Endo, 2005;Gardner et al., 2011;Marie et al., 2012), our results indicate that an ocean acidification condition of 7.6 ± 0.2 pH for 20 days has negative effects in the secretory activity of the mantle edge, affecting potentially its ability to secrete macromolecules that might be part of the organic matrix of the foliated layer in *C. gigas*.

**Figure 3.**
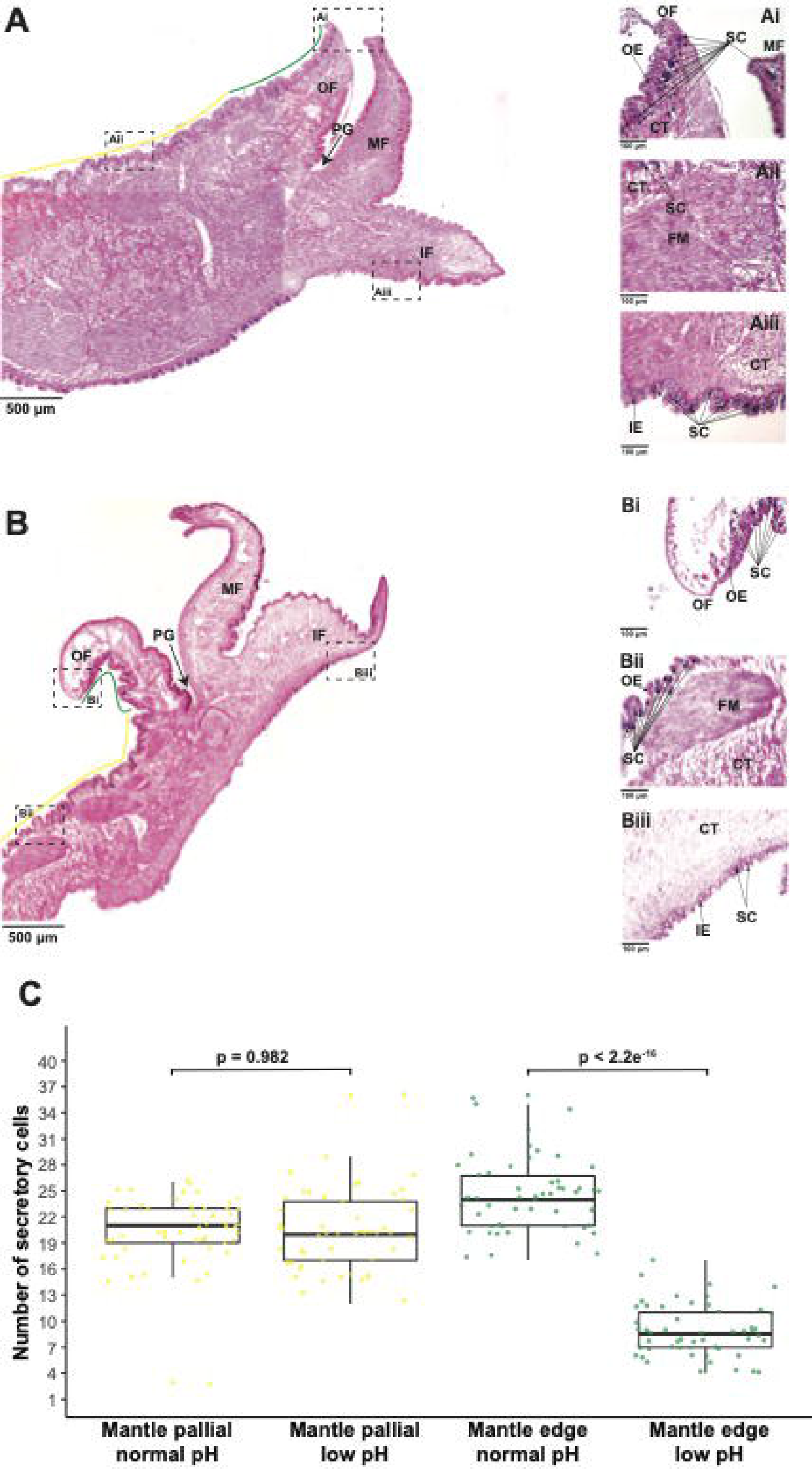
Effect of ocean acidification on the secretory activity of mantle cells. **A)** Alcian Blue-stained section through the mantle tissue of *C. gigas* kept under normal pH (8.0 ± 0.2) condition showing the mantle edge (green line) and mantle pallial (yellow line) morphogenetic zones. Scale bar = 500μm. In addition, three mantle regions are shown to indicate: **Ai)** A region of the outer fold belonging to the mantle edge zone, highlighting secretory cells in blue. Scale bar = 100μm. **Aii)** A region of the outer epithelium belonging to the mantle pallial zone, showing secretory cells in blue. Scale bar = 100μm. **Aiii)** A region of the inner fold that shows secretory cells in blue. Scale bar = 100μm. **B)** Alcian Blue-stained section through the mantle tissue of *C. gigas* kept under low pH (7.6 ± 0.2) condition showing the mantle edge (green line) and mantle pallial (yellow line) morphogenetic zones. Scale bar = 500μm. Three mantle regions are also shown to indicate: **Bi)** A region of the outer fold belonging to the mantle edge zone, highlighting secretory cells in blue. Scale bar = 100μm. **Bii)** A region of the outer epithelium belonging to the mantle pallial zone, showing secretory cells in blue. Scale bar = 100μm. **Biii)** A region of the inner fold that shows secretory cells in blue. Scale bar = 100μm. OF: outer fold. PG: periostracal groove. MF: middle fold. IF: inner fold. OE: outer epithelium. CT: connective tissue. FM: fibers muscle. IE: inner epithelium. SC: secretory cells. **C)** Number of secretory cells along the outer epithelium covering the mantle edge and mantle pallial zones in adult oysters kept under normal and low pH conditions. Yellow dots indicate secretory cells found in the mantle pallial zone, whereas green dots correspond to secretory cells in the mantle edge zone.

### Ocean acidification induces few shifts in terms of gene expression of the mantle secretome

To obtain a comprehensive snapshot of the molecular changes influenced by OA, we conducted transcriptome sequencing of adult mantle tissues from oysters exposed to normal (8.0 ± 0.2) and low (7.6 ± 0.2) pH. Raw transcriptome data are publicly available under the following NCBI BioProject PRJNA904561 and read mapping statistics to the *C. gigas* genome (Zhang et al., 2012) are shown in **Supplementary Table 1**. Genes significantly differentially expressed in mantle tissues under different pH conditions were identified using edgeR. We found 172 genes with differential expression profiles, of which 78 genes showed a down-regulation under low pH conditions, while 94 genes are over-expressed by ocean acidification (**Figure 4A and Supplementary Figure 4**). Although there are not highly represented (p < 0.05) GO terms among DEGs, most GO terms associated with up-regulated genes (>50%) belong to binding (GO:0005488), biological regulation (GO:0065007), regulation of biological processes (GO:0050789), response to stimulus (GO:0050896), and cellular process (GO:0009987) (**Supplementary Figure 5**). While most down-regulated genes (>50%) are related to the catalytic activity (GO:0003824), binding (GO:0005488), cellular process (GO:0009987), and metabolic process (GO:0008152) (**Supplementary Figure 6**). Based on the presence of conserved signal sequences, we estimate that 31 of 172 genes show differential expression between normal and decreased pH conditions (**Figure 4A**), of which 21 have associated Swissprot or PFAM annotations (**Figure 4B and Supplementary Table 2**). No specific GO term (biological process or molecular function) is over-represented in this dataset, most likely due to the low number of annotated genes. Furthermore, three DEGs encoding secreted proteins have previously been reported as SMPs in *C. gigas* (**Figure 4B and Supplementary Table 2**).

**Figure 4.**
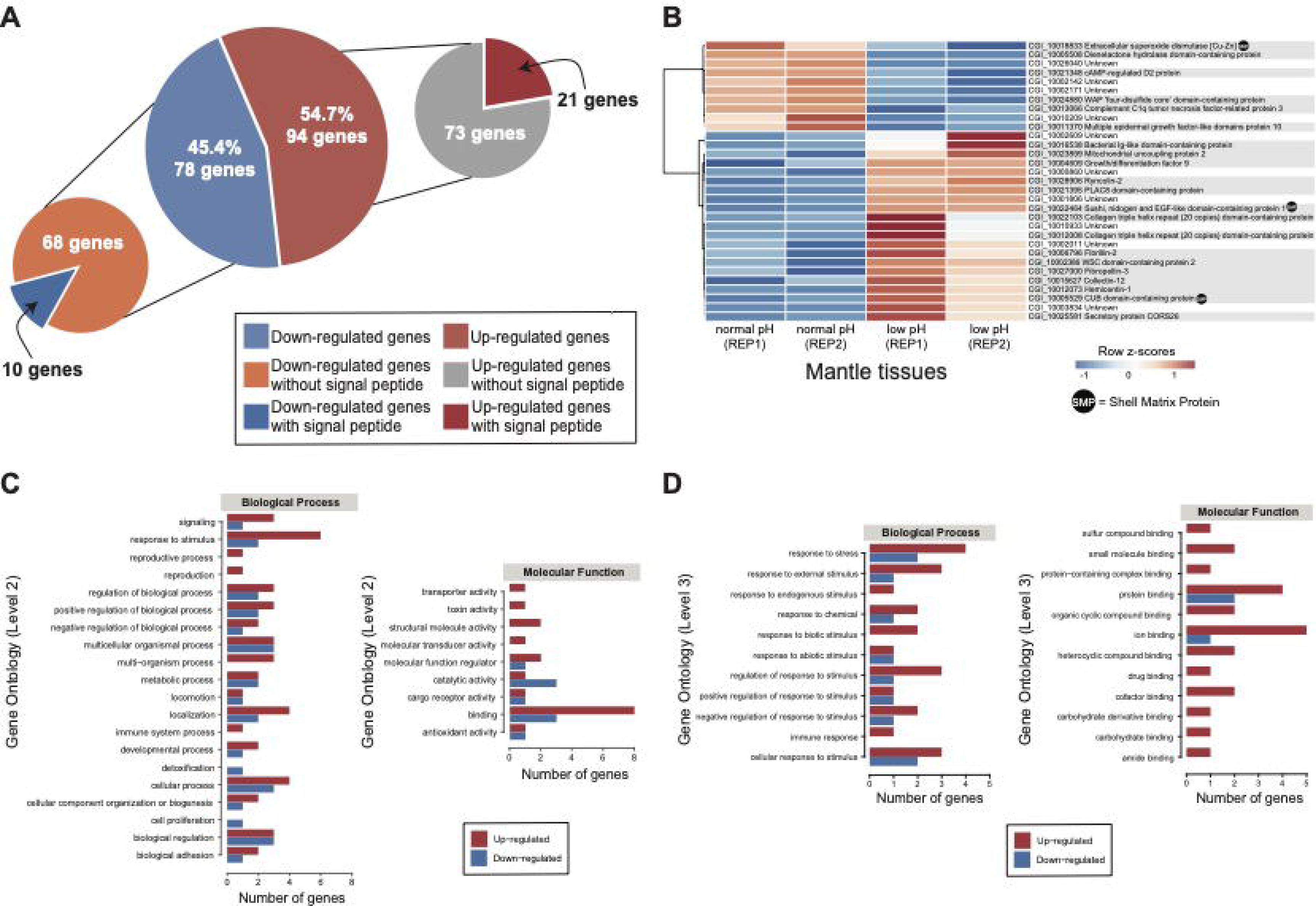
Transcriptomic response of mantle cells of oysters exposed to ocean acidification condition. **A)** A total of 172 genes were identified as differentially expressed in adult oysters reared under normal (8.0 ± 0.2) and low (7.6 ± 0.2) pH conditions and placed into two categories according to the presence or absence of a signal sequence. **B)** Heatmap of significantly differentially expressed genes encoding secreted proteins. At the bottom of each column is depicted the mantle tissue and the corresponding pH condition used to generate the RNA-Seq data. Genes are displayed as horizontal rows clustered by similar expression profiles, represented by the dendrogram to the left of the heatmap. Red indicates higher expression, while blue illustrates lower expression. Gene annotations are indicated to the right of the heatmap, and gene products (proteins) that have been previously identified as part of the organic matrix of larval/adult shells are indicated by a black dot. **C)** Distribution of differentially expressed genes encoding secreted proteins with annotated biological process GO terms (Level 2) and molecular function GO terms (Level 2). See Supplementary Table 3 for gene name information. **D)** Distribution of differentially expressed genes encoding secreted proteins with annotated biological process GO terms (Level 3) and molecular function GO terms (Level 3). See Supplementary Table 4 for gene name information.

Ten differentially expressed genes encoding secreted proteins are downregulated in the mantle tissues of oysters reared at low pH (**Figure 4B**). One DEG (Extracellular superoxide dismutase [Cu-Zn]) is an SMP and five have Swissprot or PFAM annotations (**Supplementary Table 2**). Down-regulated genes include complement C1q tumor necrosis factor-related protein 3, cAMP-regulated D2 protein, and multiple epidermal growth factor-like domains protein 10 (**Figure 4B and Supplementary Table 2**). By contrast, 21 DEGs encoding secreted proteins are upregulated in the mantle tissues of oyster exposure to low pH (**Figure 4A**), and two (Sushi, nidogen and EGF-like domain-containing protein 1 and CUB domain-containing protein) are like known SMPs (**Figure 4B and Supplementary Table 2**). Aside from SMPs, 13 genes have Swissprot or PFAM annotations (**Figure 4B and Supplementary Table 2**). Up-regulated genes include ryncolin-2, fibrillin-2, mitochondrial uncoupling protein 2, collectin-12, fibropellin-3, growth/differentiation factor 9, hemicentin-1, secretory protein CORS26, and WSC domain-containing protein 2 (**Figure 4B and Supplementary Table 2**).

The analysis of those DEGs encoding secreted proteins reveals several functional domains associated with cys-rich proteins (PLAC8 motif-binding domain), extracellular structural proteins (collagen triple helix repeat, CUB domain, F5/8 type C domain, fibrinogen beta and gamma chains C-terminal globular domain, VWA domain containing CoxE-like, von Willebrand factor type A domain, nidogen-like, AMOP domain, sushi repeat), immune proteins (bacterial Ig-like domain, C1q domain, IgGFc domain), carbohydrate-binding proteins (WSC domain, lectin C-type domain), calcium-binding proteins (complement Clr-like EGF-like, calcium-binding EGF domain, EGF domain, human growth factor-like EGF, EGF-like domain), substrate carrier proteins (mitochondrial carrier domain), chitin-binding proteins (chitin binding Peritrophin-A domain), and enzymes (dienelactone hydrolase, WAP ‘four-disulfide core’, carboxylesterase, BD-FAE, abhydrolase 3 alpha/beta hydrolase fold, copper/zinc superoxide dismutase) (**Supplementary Table 2**).

The GO analysis revealed that most DEGs (>50%) encoding secreted proteins are involved in binding (GO:0005488), response to stimulus (GO:0050896), localization (GO:0051179), cellular process (GO:0009987), and biological regulation (GO:0065007) (**Figure 4C and Supplementary Table 3**). In the binding category, six genes are likely to have protein binding function (GO:0005515) and the same number of genes seem to have ion binding (GO:0043167) function (**Figure 4D and Supplementary Table 4**). Whereas in the response to stimulus category, six genes are associated with response to stress (GO:0006950) and five genes with cellular response to stimulus (GO:0051716) (**Figure 4D and Supplementary Table 4**). Most genes related to protein binding, ion binding, response to stress, and cellular response to stimulus are upregulated in mantle cells of oysters kept at 7.6 pH (**Figures 4C and 4D**), suggesting expression shifts in these genes could be responsible for regulating pH in response to OA. Notably, three genes are found to be downregulated in mantle cells exposed to OA that exhibit catalytic functions (**Figure 4C and Supplementary Table 3**).

Four genes (fibrillin-2, secretory protein CORS26, collectin-12, and complement C1q tumor necrosis factor-related protein 3) with potential signaling functions and encoding secreted proteins are differentially expressed (**Figure 4B and Supplementary Table 2**). Three of them fibrillin-2, secretory protein CORS26, and collectin-12) are upregulated in the mantle cells of oysters reared at low pH, while complement C1q tumor necrosis factor-related protein 3 is downregulated. This result suggests a putative role of these signaling genes in the genetic regulatory network that affects OA; even though their functions are still unstudied in mollusks. No specific transcriptional regulatory genes encoding secreted proteins were found as differentially expressed (**Supplementary Table 2**).

## 4 Discussion

We conducted an integrative analysis of histological and transcriptomic data in the mantle tissue of Pacific oyster (*Crassostrea gigas*) exposure to normal (8.0) and low (7.6) pH conditions to elucidate how oysters cope with OA and examine the cellular and molecular underpinnings of this response. At the cellular level, oysters maintained in low pH conditions do not show significant differences in terms of mantle tissue anatomy, although the secretory activity of the mantle cells is diminished by OA. At the molecular level, *C. gigas* exhibited a subtle transcriptomic response to OA, and surprisingly few shifts occur in terms of gene expression.

### Effects of ocean acidification on mantle anatomy and cell morphology

Molluscan mantle epithelia have long been studied to understand the biological control of shell formation, as reviewed by Suzuki and Nagasawa (2013) and Kocot et al. (2016). Mantle tissue encloses all internal organs and is organized into different regions namely mantle edge and mantle pallial, with the mantle edge in direct contact with the prismatic shell layer and the mantle pallial with the nacreous layer (Kocot et al., 2016;Clark, 2020). Within the mantle edge, there is a series of folds with the inner and middle folds involved in water inflow and sensory functions, respectively, whilst the outer fold is involved in shell secretion (Clark, 2020;Clark et al., 2020). Most bivalves have three mantle folds (Suzuki and Nagasawa, 2013;Clark et al., 2020), although there are exceptions where the outer fold is duplicated (Waller, 1980) or the inner fold is fused (Sleight et al., 2016). In bivalves, the shell varies greatly in terms of shape, color, size, and thickness, which might be related to the different morphologies of the mantle tissue (Parvizi et al., 2017). The mantle of *C. gigas* possesses many similarities to that of pearl oysters (*Pinctada* spp.), including a single periostracal groove, three folds in the mantle edge, and an outer fold epithelium encompasses mainly columnar epithelial cells (Dix, 1972;Jabbour-Zahab et al., 1992;Parvizi et al., 2017;Zhu et al., 2021;Han et al., 2022). However, the structure of the periostracal groove is somehow different in that two cell types are distinguishable in *C. gigas*, columnar cells to the base and planar cells to the tip of the periostracal groove, which are not observed in pearl oysters (Jabbour-Zahab et al., 1992;Parvizi et al., 2017). Given the morphological similarities in the mantle tissue between *C. gigas* and pearl oysters, it is likely that the differences observed, in terms of the shape and microstructure of their shells (Farre et al., 2011;Lee et al., 2011;Marie et al., 2011;Marie et al., 2012;Mouchi et al., 2016;Checa et al., 2018;Deng et al., 2022), are the product of different shell matrix proteins secreted by the mantle in a species-specific manner (Aguilera et al., 2017), rather than as a product of mantle morphology. Inside the mantle tissue, there are contractile muscle fibers possibly involved in mantle retraction and adipose tissue containing high glycogen content (Berthelin et al., 2000;Álvarez Nogal and Molist García, 2015;Parvizi et al., 2017). It is well known that shell formation is a high-demand energy process (Palmer, 1992;Waldbusser et al., 2013;Clark et al., 2020); therefore, the use of glycogen as an energy resource is no doubt an economic method employed by mollusks to construct their shells.

Most studies evaluating the effect of OA in adult individuals have reported negative outcomes, including reduced calcification rates as well as decreased shell growth and strength, but also increased shell dissolution, reviewed by Parker et al. (2013). However, the impact of OA in terms of mantle morphology is scarce (Wei et al., 2015). We here found that the effect of reduced pH in the mantle tissue of *C. gigas* for 20 days was undetectable with no apparent detrimental effects on cell morphology between hypercapnic and normocapnic oysters, which is in concordance with a previous study exposing adult oysters to 28 days (Wei et al., 2015). This might indicate that hypercapnic adult individuals develop physiological strategies to allocate extra energy for pH homeostasis under a decreased pH external environment. However, this mechanism remains speculative and requires further investigation.

### Ocean acidification provokes a reduced secretory activity of mantle edge cells

Ocean acidification (OA) triggered by anthropogenic activities is accompanied by an increase of CO_2_, which when dissolved in water becomes carbonic acid (H_2_CO_3_) (Falkowski et al., 2000;Caldeira and Wickett, 2003;Doney et al., 2009). This carbonic acid dissociates into bicarbonate (HCO_3_^−^) and a hydrogen ion (H^+^), the latter of which makes the water more acidic (Doney et al., 2009;Hurd et al., 2019). The fabrication of the mollusk shell involves the selective uptake of calcium (Ca^2+^) and bicarbonate (HCO^3−^) ions from the environment that are eventually transported through the outer epithelium of the mantle into the mineralization front (Jodrey, 1953;Sillanpää et al., 2016;Sillanpää et al., 2018;Sillanpää et al., 2020). The HCO^3-^ is used as the carbon source by mollusks to produce the shell through a process during which another hydrogen ion (H^+^) is released, and thus consequently alters the acid-base homeostasis of the mantle cells (Ramesh et al., 2020). Therefore, it is essential to understand the cellular response and secretory activity employed by mantle cells to cope with OA.

Most mollusks possess a multi-layered shell that is formed by specialized secretion of distinct morphogenetic zones of the mantle (Kocot et al., 2016). The outermost layer, known as the periostracum, comprises proteins secreted from the periostracal groove (Zhang et al., 2006;Checa and Harper, 2010;Nakayama et al., 2013). The middle prismatic layer is thought to be controlled by genes expressed in the mantle edge region, whereas the inner nacreous layer is likely controlled by genes expressed in the mantle pallial (Takeuchi and Endo, 2005;Gardner et al., 2011;Marie et al., 2012;Herlitze et al., 2018). This spatial dichotomy in expression is corroborated by ultrastructural investigations of the mantle epithelium, indicating cellular differentiation of prism- and nacre-secreting mantle cells (Fang et al., 2008). In pearl oysters, two major secretory cells have been described – basophilic and acidophilic – in the mantle, where the former type is characterized by sulfated acidic mucosubstances (Parvizi et al., 2017). We found that most of the mucous cells in the mantle stained blue with the combined technique of AB-PAS showing that they contain acidic mucosubstances. These AB-PAS positive cells are distributed along the outer mantle epithelium, covering the mantle edge and mantle pallial zones. These acidic mucosubstances might presumably have an active role in the mineralization of the shell by aiding with the absorption or binding of calcium ions (Breedham, 1958;Kapur and Gibson, 1967;Álvarez Nogal and Molist García, 2015). The adult shell of *C. gigas* is mostly composed of calcite and a small portion of aragonite in the myostracum (Lee et al., 2011;Marie et al., 2011;Mouchi et al., 2016;Checa et al., 2018). In contrast to most bivalves, *C. gigas* shell is made up of a shell-like foliae layer and lenses of chalk (Marie et al., 2011;Checa et al., 2018), a highly porous and poorly organized microstructure that is only found in Ostreidae (Mouchi et al., 2016;Checa et al., 2018). Recent structural and crystallographic observations have depicted that growth lines are continuous between the foliae and chalk, suggesting both layers are made simultaneously by different zones of the mantle tissue, where the foliated layer is made by the mantle edge and the chalky layer by the mantle pallial (Banker and Sumner, 2020). In oysters kept in low pH conditions, the outer epithelium of the mantle edge shows fewer AB-PAS positive cells compared with the mantle pallial zone, suggesting a decrease in the secretory activity in mantle cells underlying the foliated layer that possibly affects its formation.

### Transcriptome analysis provides new insights into the secretory response of adult oysters against ocean acidification

Transcriptional profiling has been performed to provide a more detailed picture of the molecular events underpinning the physiological response of marine animals to OA (Strader et al., 2020). Here, we found a relatively low number of genes (172) that were significantly differentially expressed under OA, suggesting a subtle transcriptomic response to OA in *C. gigas* mantle cells after 20 days of low-pH exposure.

Gene ontology classification of these DEGs showed that low pH (7.6) triggers the regulation of cellular and biological processes but prominently suppresses some metabolic processes and catalytic activities. These results were consistent with previous studies on *C. gigas* larvae at 14-16 hours post-fertilization exposed to similar levels of low-pH stress (de Wit et al., 2018). In addition, we found that response to stimulus and immune-related genes are up-regulated in mantle tissues of oysters in the low pH group. Immune-related genes have been found down-regulated in the hemocytes of adult *C. gigas*, as well as in *C. gigas* larvae, exposed to OA (Wang et al., 2016;Wang et al., 2020;Dineshram et al., 2021), but it has also been seen that immune-related genes are up-regulated in other oysters (Goncalves et al., 2016;Goncalves et al., 2017). We also found that catalytic and oxidoreductase genes were downregulated in oysters kept in low pH conditions, which agrees with what happens before larval shell formation in *C. gigas* (Liu et al., 2020); but it is inconsistent with what has been observed in the adult mantle tissue of *C. virginica* (Downey-Wall et al., 2020). In contrast to previous studies that showed higher expression of genes involved in metabolism and oxidative stress under OA in either *C. gigas* larvae or hemolymph from adult oysters (Wang et al., 2016;Cao et al., 2018;Dineshram et al., 2021), we did not find such changes in mantle cells of *C. gigas* after exposure to low pH during 20 days of exposure. Together, these data, and previous studies in other marine metazoans (Strader et al., 2020), suggest that marine organisms vary dramatically in their transcriptomic responses to low pH, with differences among closely related animal groups and even between life-history stages.

To examine the effects of OA on the mantle secretory activity at the molecular level, we focused on DEGs encoding secreted proteins – the so-called mantle secretome (Jackson et al., 2006;Kocot et al., 2016;Aguilera et al., 2017) – because those proteins are likely to be embedded within the shell. Among genes encoding secreted proteins that were significantly differentially expressed, we found the presence of domains of diverse nature that cover a large set of functions. Some are collagen-like, von Willebrand factor type A, and sushi-like domains that are associated with extracellular matrix binding molecules (Marin, 2020). Von Willebrand type A domains are usually found in most - if not all - molluscan organic matrices and exhibit an adhesion function through protein-protein interactions, playing a role in 3D framework structuring (Whittaker and Hynes, 2002;Arivalagan et al., 2017;Marin, 2020). Saccharide domains comprise the WSC and lectin C-type domains. Many molluscan shell-associated proteins have lectins that are calcium-dependent but do not have a structural role as framework constituents; indeed, they may exert a function in immunity and protection (Marin, 2020). Calcium-binding domains include several kinds of EGF domains. These domains are involved in a diverse array of functions, including intracellular signaling mediated by calcium (Wouters et al., 2005). Notably, the EGF-like domain has been reported in tandem repeats in *Pinctada* and *Crassostrea* shells (Marie et al., 2011;Marie et al., 2012), and this repeating arrangement is favored during evolution due to its stability against enzyme attack through the formation of compact dimeric or tetrameric pseudo-symmetrical structures (Arivalagan et al., 2017). Domains related to immunity and protective mechanisms comprise C1q and WAP ‘four-disulfide core’ domains. The C1q domain that belongs to the C1q protein family (Ghebrehiwet et al., 2012) has been previously identified in the *C. gigas* shell (Arivalagan et al., 2017). WAPs are highly conserved proteinase inhibitor domains found in vertebrate and invertebrate species (Smith, 2011). WAP domain-containing proteins play major roles in gastropod shell formation (Shen et al., 1997;Treccani et al., 2006;Marie et al., 2010), inhibiting calcium carbonate crystal growth *in vitro* (Treccani et al., 2006). However, the function of WAP domain-containing proteins in *Crassostrea* is unknown. Most secreted genes annotated as downregulated do not possess similarity with previously described Swissprot sequences, even though some of them have functional domains that give some glimpse into their putative role. Aside from unknown genes, other down-regulated genes exhibited similarity to complement C1q tumor necrosis factor-related proteins; multiple growth factor-like domains protein 10, and metabolic proteins such as cAMP-regulated D2 gene and extracellular superoxide dismutase [Cu-Zn]. Regarding annotated upregulated genes encoding secreted proteins, these include ryncolin-2, fibrillin-2, mitochondrial uncoupling protein 2, collectin-12, fibropellin-3, growth/differentiation factor 9, hemicentin-1, secretory protein CORS26, and WSC domain-containing protein 2. Although the function of most - if not all - of these genes is unknown in mollusks, there are some upregulated DEGs encoding secreted proteins exhibiting similarity with genes that have been implicated in biomineralization in other species, for example, secretory protein CORS26 (Maeda et al., 2001), fibrillin-2 (Arteaga-Solis et al., 2011;Mead et al., 2022), and hemicentin-1 (Ramos-Silva et al., 2013;Germer et al., 2015;Jackson et al., 2015). Indeed, hemicentin-1, a multimodular extracellular matrix protein that interacts with basement membrane proteins to confer tissue linkage (Zhang et al., 2022), has been reported as downregulated in *C. virginica* larvae exposed to low pH (Barbosa et al., 2022), but also as upregulated in *C. gigas* larvae and adults exposed to low pH (Timmins-Schiffman et al., 2014;Dineshram et al., 2021), as in this study as well. Furthermore, secreted proteins involved in protein binding, ion binding, response to stress, and cellular response to stimulus were upregulated in mantle tissues of oysters kept at 7.6 pH. However, a previous study on *C. gigas* larvae has shown a suppression of proteins involved in response to stimulus at a 7.4 pH (Dineshram et al., 2016). Our findings are not in line with those of Dineshram et al. (2016) and strongly suggest that the effects of OA vary considerably between different life-story stages in the same species. Within ion binding GO terms, calcium ion binding genes (fibrillin-2, hemicentin-1, and sushi, nidogen and EGF-like domain-containing protein 1) were positively regulated by OA. The stimulation of calcium-related genes was shown in *C. gigas* larvae to survive and develop under OA but at a slower rate (de Wit et al., 2018). In a previous study in adult *C. gigas* but focusing on the hemocytes response, a series of calcium-binding genes and calcium signaling pathways were also upregulated in response to OA indicating the regulation of calcium homeostasis under low pH (Wang et al., 2020). By contrast, calcium-dependent signaling pathways were suppressed in the pearl oyster *Pinctada fucata* under OA (Li et al., 2016) but whereas in the case of *C. virginica*, no significant difference in terms of calcium-related gene expression was observed upon exposure to low pH (Downey-Wall et al., 2020). Thus, mantle cells likely activate different cellular stress responses and modulate genes involved in calcium metabolism and pathways to cope with low pH. Some secreted genes encoding shell matrix proteins embedded in the calcified adult shell of *C. gigas* (Zhang et al., 2012) were also found to be significantly differentially expressed between hypercapnic and normocapnic oysters, suggesting that physical or mechanical properties of the shell might be affected to some degree by OA.

Altogether, we show that the transcriptome response of mantle cells under OA condition for 20 days is subtle, but there are significant changes in metabolic genes, shell-building genes, and calcium-dependent pathways.

## 5 Conclusion

We evaluated the effects of low pH in terms of cell morphology, secretory activity, and gene expression profiles in the mantle cells of adult oysters exposed to OA conditions for 20 days. During this time, we found that the morphology of mantle cells is not affected by exposure to OA.

Nevertheless, mantle cell secretory activity is influenced by OA but in a mantle zone specific manner. It appears likely that mantle edge cells are more susceptible to OA. The modular nature of the mantle tissue (Herlitze et al., 2018), along with the difference in secretory activity in mantle edge cells, suggests that the foliated shell layer might be the first compromised due to OA.

Furthermore, our study demonstrated a subtle transcriptomic response of adult mantle cells with few shifts in gene expression profiles. However, we identified several OA-responsive genes related to cellular and biological processes involved in the metabolic process, catalytic activity, and ion binding. Once we focused on DEGs encoding secreted proteins, mantle cells differentially expressed genes involved in metabolic and calcium-dependent pathways and shell biomineralization. Since most OA studies have been conducted on whole embryos or non-calcifying tissues (Leung et al., 2022), this study provides valuable morphological, secretory, and transcriptomic information on the mantle tissue for understanding the capacity of *C. gigas* to respond to OA.

## 7 Author Contributions

FA conceived the study, conducted bioinformatics and data analysis, and wrote the manuscript with contributions from all other authors. NZ-S performed fertilization experiments under normal and low pH conditions with assistance from FA and conducted the histological analysis. IP-B performed histological analysis. CQ performed bioinformatic analysis. All authors have read and approved the final manuscript.

## 9 Acknowledgments

We acknowledge all members from the Group of Developmental Processes (GDeP) at the University of Concepcion for fruitful discussions on this project. We would also like to thank anonymous reviewers for their insightful comments that improved our manuscript.

## 11 Supplementary Material

The Supplementary Material for this article can be found online at

**Supplementary Figure 1 |** Toluidine blue-stained section through the mantle tissue of *C. gigas* kept under normal pH (8.0 ± 0.2) condition (PDF).

**Supplementary Figure 2 |** Hematoxylin and Eosin-stained sections through the mantle tissue of *C. gigas* kept under low pH (7.6 ± 0.2) condition (PDF).

**Supplementary Figure 3 |** Toluidine blue-stained sections through the mantle tissue of *C. gigas* kept under low pH (7.6 ± 0.2) condition (PDF).

**Supplementary Figure 4 |** The volcano plow shows significantly differentially expressed genes in mantle cells of oysters kept under normal (8.0 ± 0.2) and low pH (7.6 ± 0.2) conditions (PDF).

**Supplementary Figure 5 |** Gene Ontology (GO) classification of up-regulated genes in mantle cells exposed to ocean acidification conditions (pH = 7.6 ± 0.2) (PDF).

**Supplementary Figure 6 |** Gene Ontology (GO) classification of down-regulated genes in mantle cells exposed to ocean acidification conditions (pH = 7.6 ± 0.2) (PDF).

**Supplementary Table 1 |** Per sample mapping summary for adult mantle RNA-Seq data.

**Supplementary Table 2 |** Genes differentially expressed (and encoding secreted proteins) in mantle cells of oysters kept under normal (8.0 ± 0.2) and low (7.6 ± 0.2) pH conditions.

**Supplementary Table 3 |** Gene Ontology terms (Level 2) of differentially expressed genes encoding secreted proteins in the mantle cells of oysters kept under normal (8.0 ± 0.2) and low (7.6 ± 0.2) pH conditions.

**Supplementary Table 4 |** Gene Ontology terms (Level 3) of differentially expressed genes encoding secreted proteins in the mantle cells of oysters kept under normal (8.0 ± 0.2) and low (7.6 ± 0.2) pH conditions.

## 1 Data Availability Statement

Raw transcriptomic reads have been deposited in NCBI under Bioproject accession number: PRJNA904561.

