## SupplementaryFiles for "Secretory and transcriptomic responses of mantle cells to low pH in the Pacific oyster (*Crassostrea gigas*)"

Supplementary Material

### 1 Supplementary Data

Supplementary Material should be uploaded separately on submission. Please include any supplementary data, figures and/or tables. All supplementary files are deposited to FigShare for permanent storage and receive a DOI.

Supplementary material is not typeset so please ensure that all information is clearly presented, the appropriate caption is included in the file and not in the manuscript, and that the style conforms to the rest of the article. To avoid discrepancies between the published article and the supplementary material, please do not add the title, author list, affiliations or correspondence in the supplementary files.

### 2 Supplementary Figures and Tables

#### 2.1 Supplementary Figures

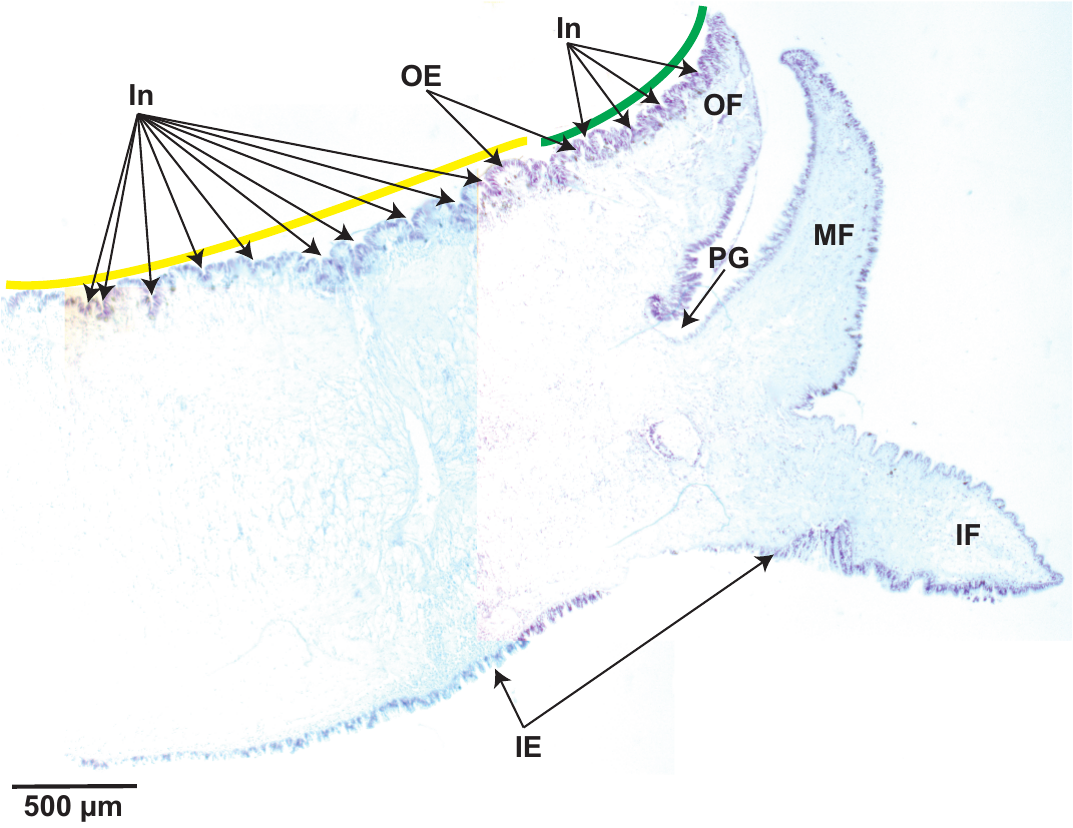

**Supplementary Figure 1.** Toluidine blue-stained section through the mantle tissue of *C. gigas* kept under normal pH (8.0 ± 0.2) condition. The green line is highlighting the mantle edge, while the yellow line is showing the mantle pallial. OE: outer epithelium. OF: outer fold. PG: periostracal groove. MF: middle fold. IF: inner fold. IE: inner epithelium. In: Invaginations of the mantle tissue along the mantle edge and mantle pallial zones. Scale bar = 500µm.

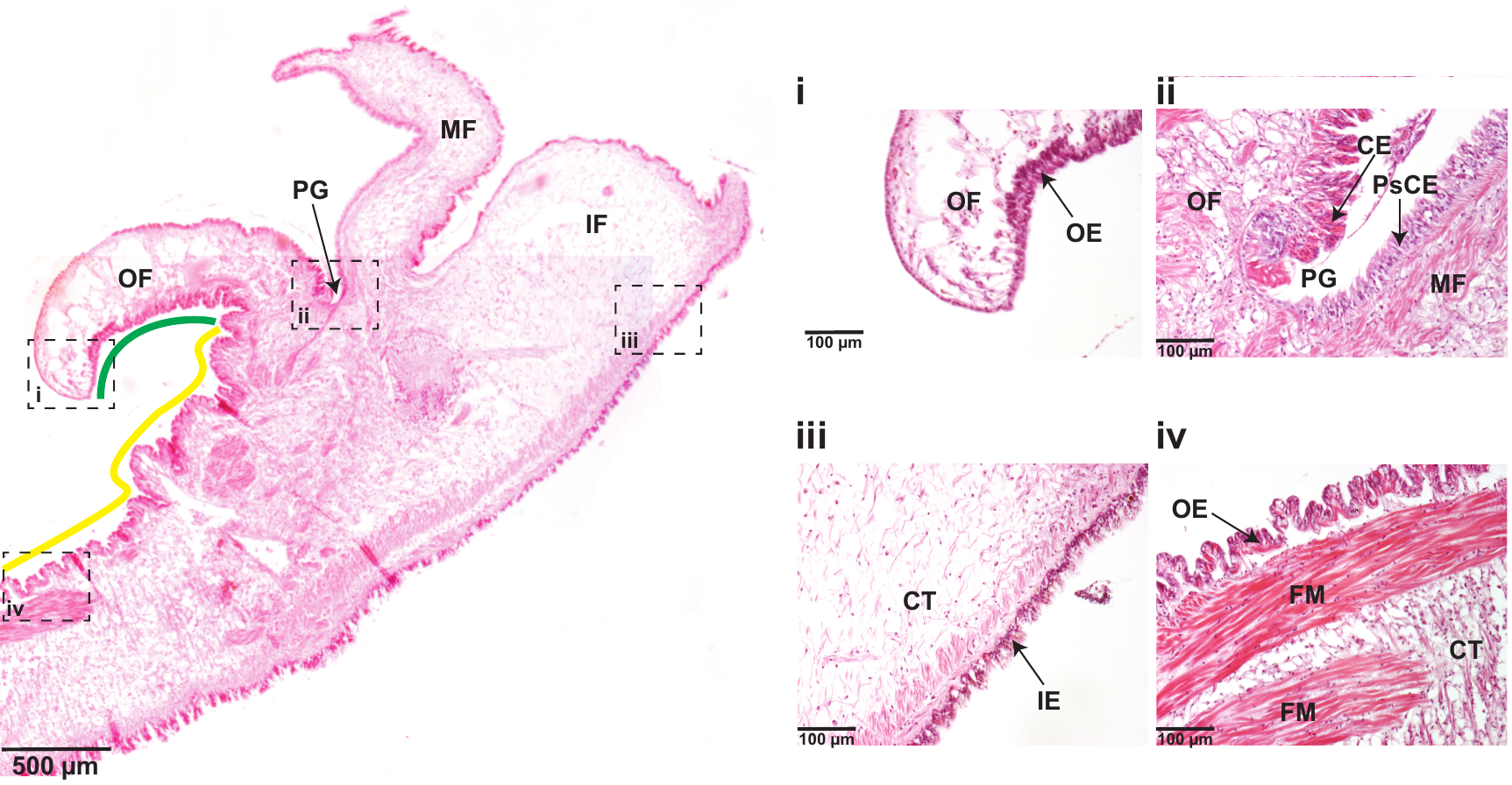

**Supplementary Figure 2.** Hematoxylin and Eosin-stained sections through the mantle tissue of *C. gigas* kept under low pH (7.6 ± 0.2) condition. An overview of mantle tissue anatomy depicting the mantle edge (green line) and mantle pallial (yellow line) zones. Scale bar = 500µm. **i)** A region of the outer fold showing the outer epithelium. Scale bar = 100µm. **ii)** Base of the periostracal groove depicting the columnar epithelium on the outer fold side and the pseudostratified columnar epithelium, with nuclei at different heights, on the middle fold side. Scale bar = 100µm. **iii)** Transverse section through the inner fold showing the inner epithelium. Scale bar = 100µm. **iv)** A region of the mantle pallial zone depicting cells of the outer epithelium and fibers muscle. Scale bar = 100µm. OF: outer fold. PG: periostracal groove. MF: middle fold. IF: inner fold. OE: outer epithelium. CE: columnar epithelium. PsCE: pseudostratified columnar epithelium. CT: connective tissue. FM: fibers muscle.

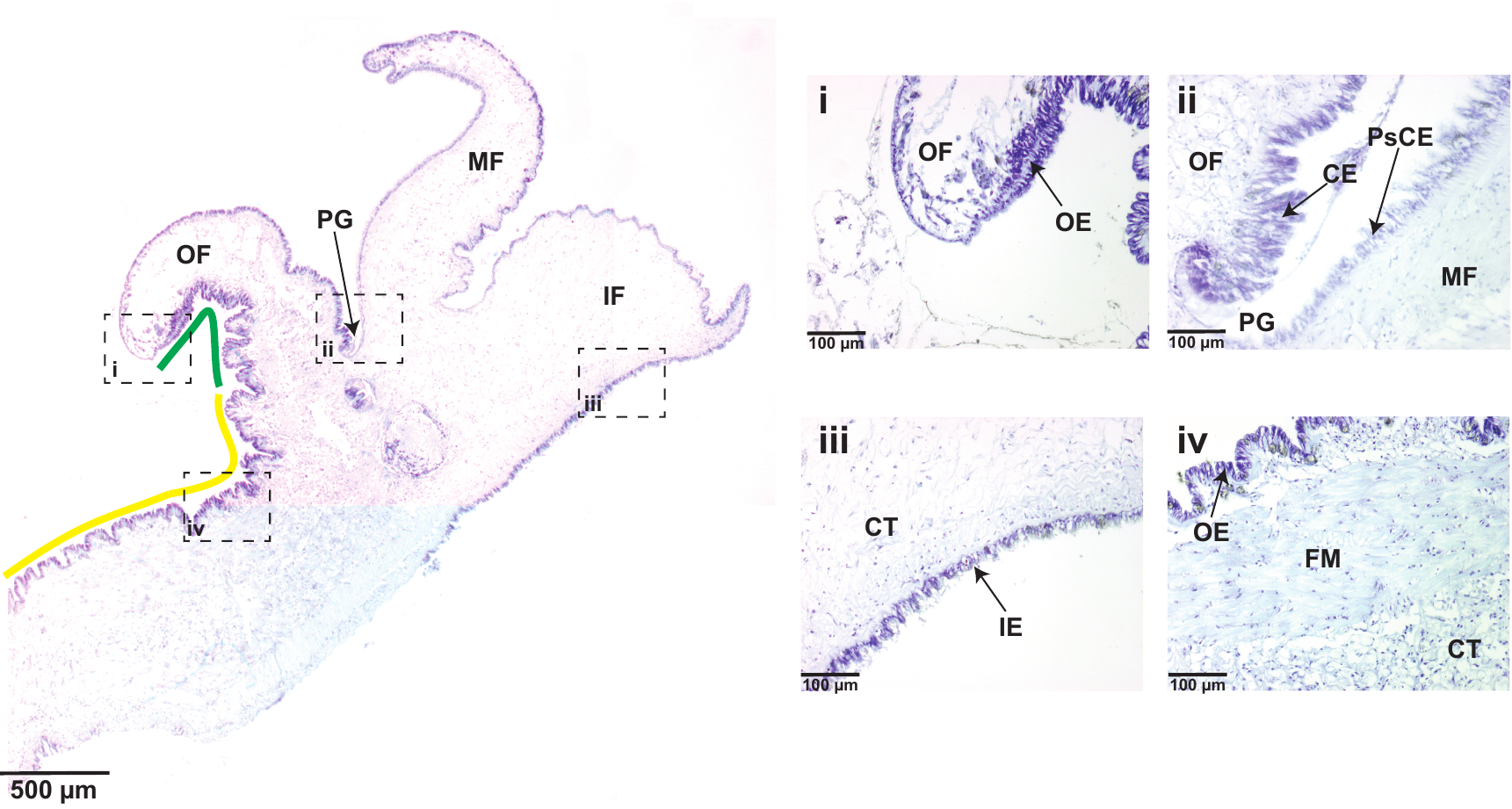

**Supplementary Figure 3.** Toluidine blue-stained sections through the mantle tissue of *C. gigas* kept under low pH (7.6 ± 0.2) condition. An overview of mantle tissue anatomy depicting the mantle edge (green line) and mantle pallial (yellow line) zones. Scale bar = 500µm. **i)** A region of the outer fold showing the outer epithelium. Scale bar = 100µm. **ii)** Base of the periostracal groove depicting the columnar epithelium on the outer fold side and the pseudostratified columnar epithelium, with nuclei at different heights, on the middle fold side. Scale bar = 100µm. **iii)** Transverse section through the inner fold showing the inner epithelium. Scale bar = 100µm. **iv)** A region of the mantle pallial zone depicting cells of the outer epithelium and fibers muscle. Scale bar = 100µm. OF: outer fold. PG: periostracal groove. MF: middle fold. IF: inner fold. OE: outer epithelium. CE: columnar epithelium. PsCE: pseudostratified columnar epithelium. CT: connective tissue. FM: fibers muscle.

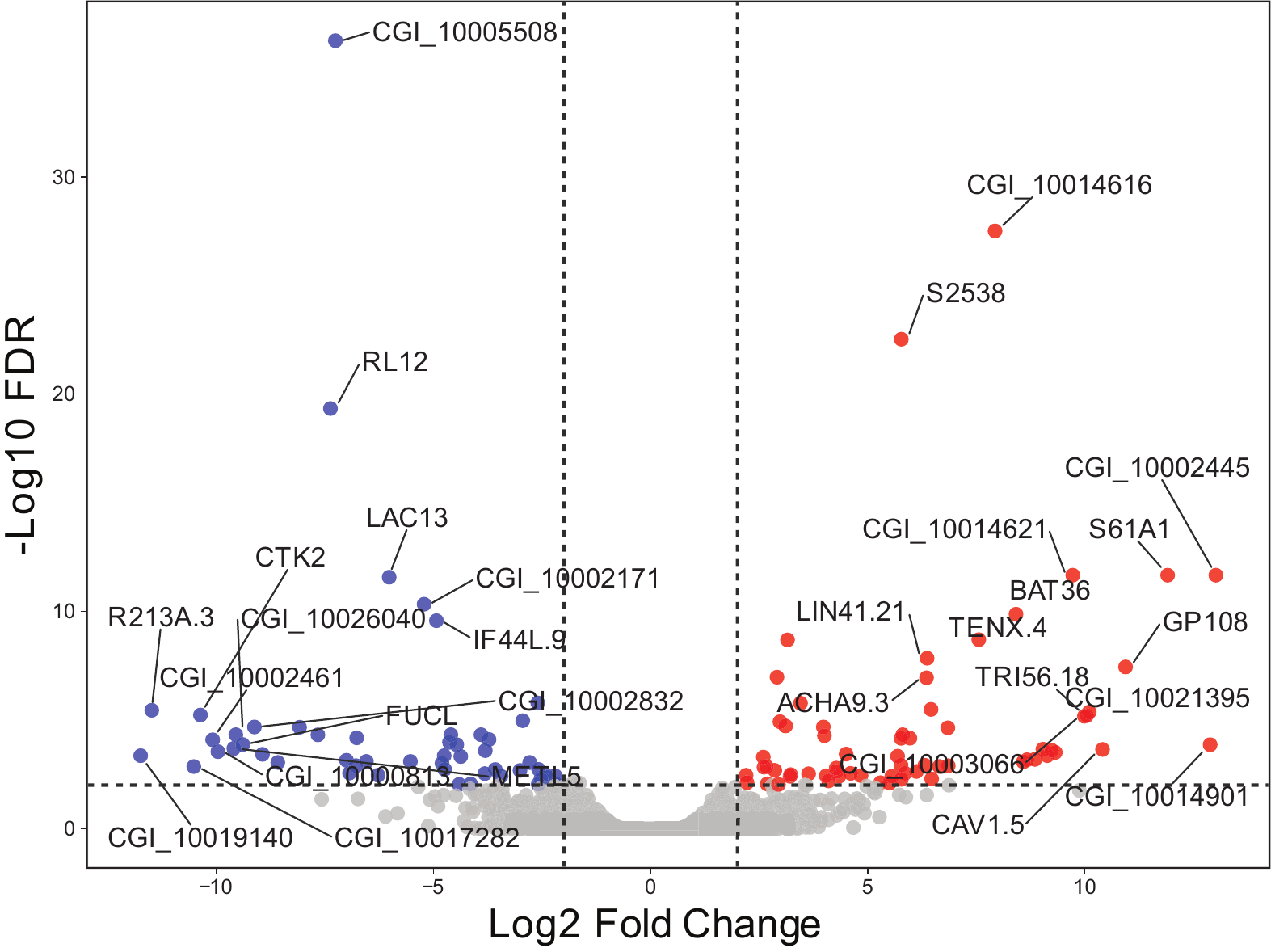

**Supplementary Figure 4.** The volcano plot shows significantly differentially expressed genes in mantle cells of oysters kept under normal (8.0 ± 0.2) and low pH (7.6 ± 0.2) conditions. The x-axis represents log2-transformed fold changes, while the y-axis represents -log10 (FDR). FRD = False Discovery Rate. Dashed lines indicate the thresholds used to define a gene as differentially expressed. Red dots depict up-regulated genes, whereas blue dots represent down-regulated genes. Gray dots show genes that are not statistically significant. Some up- and down-regulated genes are highlighted.

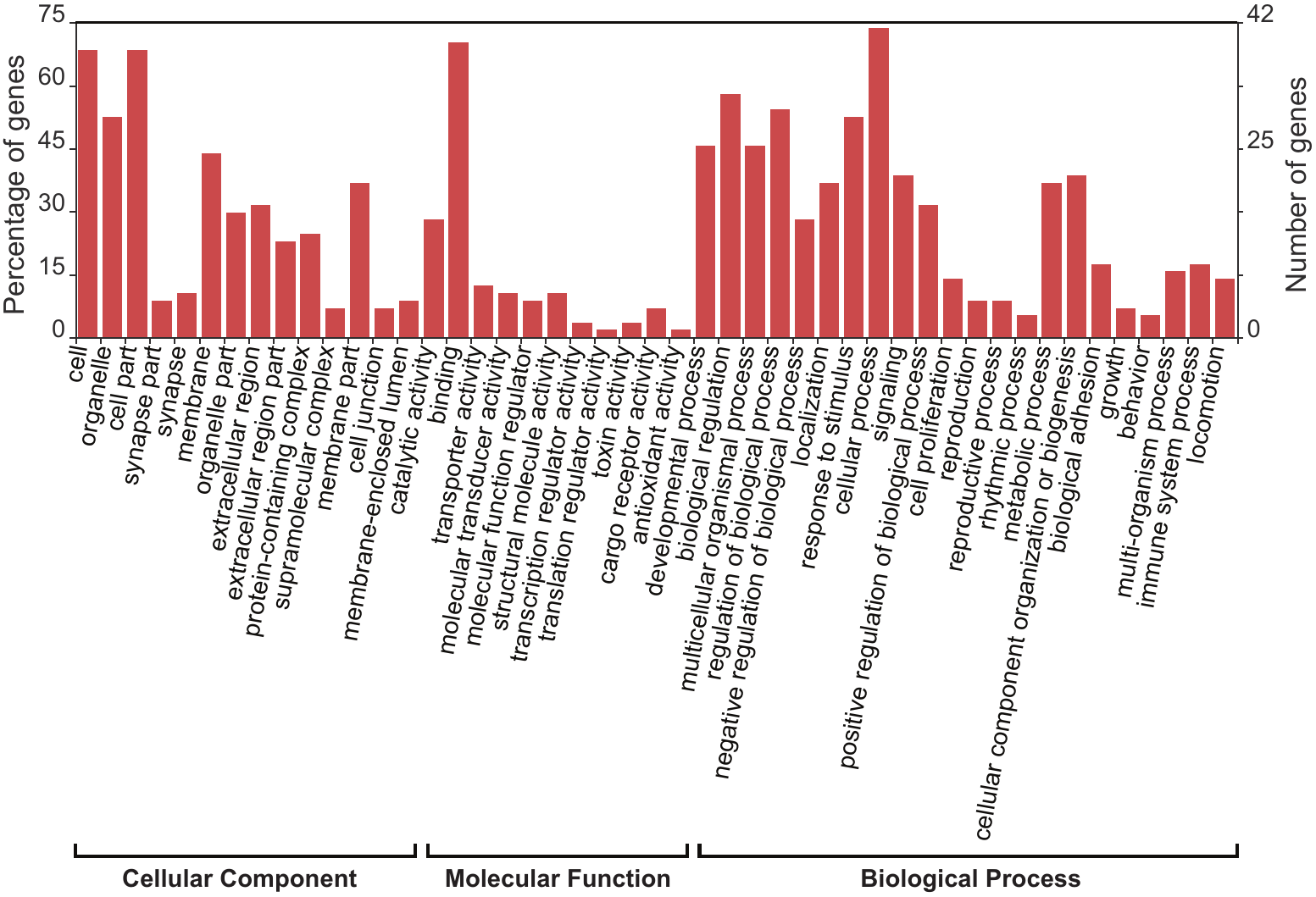

**Supplementary Figure 5.** Gene Ontology (GO) classification of up-regulated genes in mantle cells exposed to ocean acidification conditions (pH = 7.6 ± 0.2). Histogram showing summary for the three main GO categories: cellular compartment, molecular function, and biological process. The left axis indicates the percentage of sequences in each category, whereas the right axis shows the total number of genes in each category.

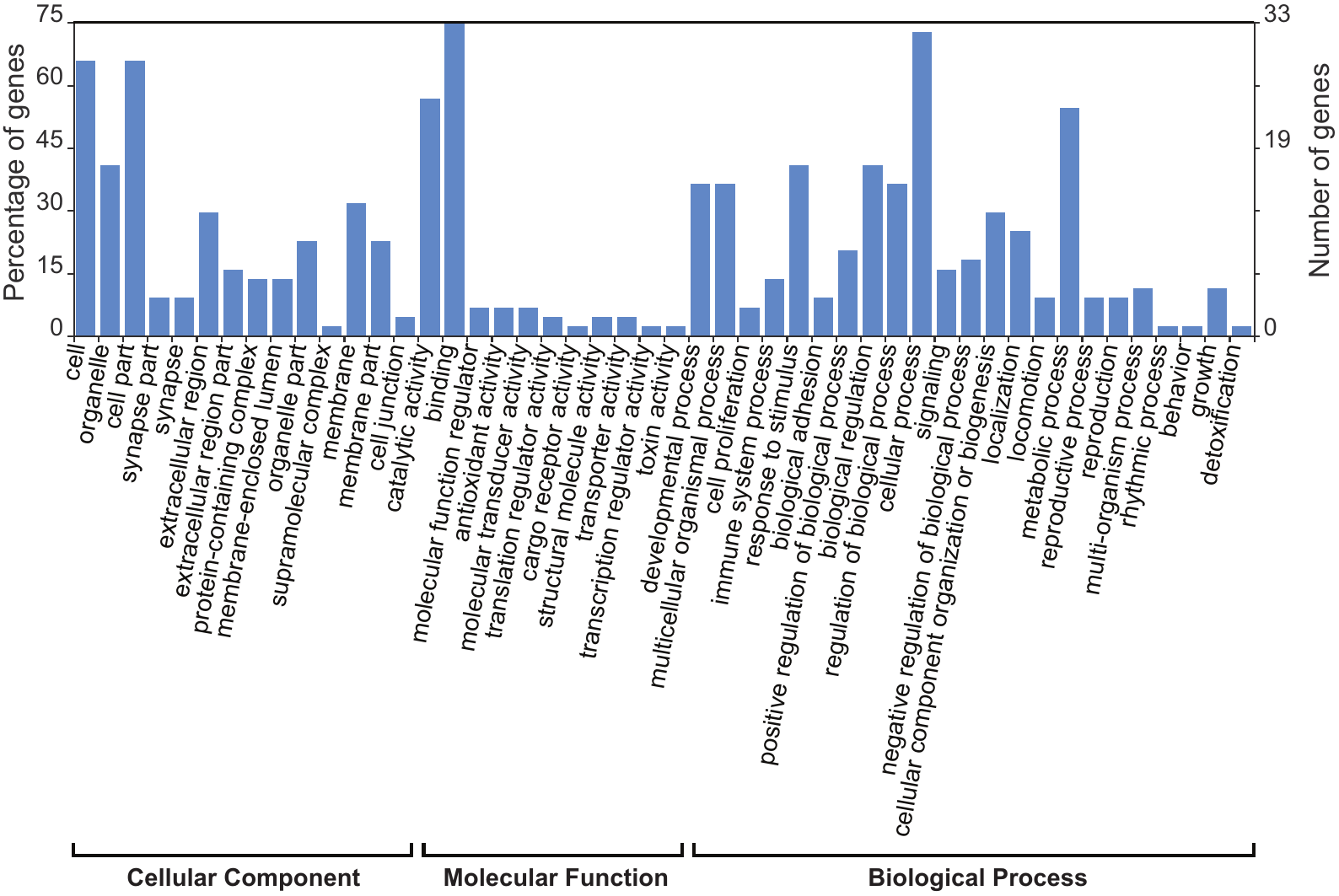

**Supplementary Figure 6.** Gene Ontology (GO) classification of down-regulated genes in mantle cells exposed to ocean acidification conditions (pH = 7.6 ± 0.2). Histogram showing summary for the three main GO categories: cellular compartment, molecular function, and biological process. The left axis indicates the percentage of sequences in each category, whereas the right axis shows the total number of genes in each category.

#### 2.3 Supplementary Tables

**Supplementary Table 1.** Per sample mapping summary for adult mantle RNA-Seq data.

|  | **Normal pH (REP1)** | **Normal pH (REP2)** | **Low pH (REP1)** | **Low pH (REP2)** |
| --- | --- | --- | --- | --- |
| Sequencing reads (paired-end) | 44,405,304 | 40,509,332 | 39,941,065 | 40,942,544 |
| Uniquely mapped reads | 34,656,799 | 31,213,085 | 29,810,369 | 30,875,693 |
| Percentage of uniquely mapped reads | 78.05% | 77.05% | 74.64% | 75.41% |
| Multi-mapped reads | 6,493,071 | 5,432,783 | 7,431,055 | 7,306,197 |
| Percentage of multi-mapped reads | 14.63% | 13.41% | 18.61% | 17.85% |
| Percentage of unmapped reads | 7.33% | 9.54% | 6.76% | 6.74% |

**Supplementary Table 2.** Genes differentially expressed (and encoding secreted proteins) in mantle cells of oysters kept under normal (8.0 ± 0.2) and low (7.6 ± 0.2) pH conditions.

| **Gene ID** | **Log2 Fold change** | **FDR** | **Swissprot top BLAST hit** | **PFAM domains** | **Shell matrix protein reciprocal hit?** | **Likely signaling function?** | **Likely transcription factor?** |
| --- | --- | --- | --- | --- | --- | --- | --- |
| CGI_10021395 | 10.1033453 | 4.34670574e-06 | No hit but it has PFAM domain | PLAC8 family (PF04749) | No | No | No |
| CGI_10012008 | 9.87438835 | 0.017484751172 | No hit but it has PFAM domain | Collagen triple helix repeat (20 copies) (PF01391) | No | No | No |
| CGI_10003834 | 9.13776056 | 0.000439028973 | Unknown | - | No | No | No |
| CGI_10022103 | 6.87029233 | 0.010086156939 | No hit but it has PFAM domain | Collagen triple helix repeat (20 copies) (PF01391) | No | No | No |
| CGI_10010933 | 6.65019318 | 0.001386118863 | Unknown | - | No | No | No |
| CGI_10012073 | 5.50350886 | 0.007972336039 | Hemicentin-1 | - | No | No | No |
| CGI_10016538 | 5.16694363 | 0.023359784931 | No hit but it has PFAM domain | Bacterial Ig-like domain (PF12245) | No | No | No |
| CGI_10025581 | 4.97561528 | 0.011147513297 | Secretory protein CORS26 | C1q domain (PF00386) | No | Yes | No |
| CGI_10002609 | 4.80429357 | 0.048402077422 | Unknown | - | No | No | No |
| CGI_10002386 | 4.74556569 | 0.041357496199 | WSC domain-containing protein 2 | WSC domain (PF01822) | No | No | No |
| CGI_10002011 | 4.50359686 | 0.000371908715 | Unknown | - | No | No | No |
| CGI_10005529 | 4.04333978 | 0.003685891296 | No hit but it has PFAM domain | CUB domain (PF00431); F5/8 type C domain (PF00754) | Yes (Zhang et al. 2012) | No | No |
| CGI_10028906 | 3.58818609 | 0.010501236731 | Ryncolin-2 | Fibrinogen beta and gamma chains, C-terminal globular domain (PF00147) | No | No | No |
| CGI_10006796 | 3.42216791 | 0.042965629247 | Fibrillin-2 | Complement Clr-like EGF-like (PF12662); Calcium-binding EGF domain (PF07645); EGF domain (PF12947); Human growth factor-like EGF (PF12661); IgGFc binding protein (PF17517) | No | Yes | No |
| CGI_10023899 | 3.34339390 | 0.042796799005 | Mitochondrial uncoupling protein 2 | Mitochondrial carrier protein (PF00153) | No | No | No |
| CGI_10001806 | 2.97893379 | 1.20150319e-05 | Unknown | - | No | No | No |
| CGI_10015627 | 2.80472855 | 0.010302798625 | Collectin-12 | Lectin C-type domain (PF00059) | No | Yes | No |
| CGI_10027000 | 2.66872469 | 0.001391876578 | Fibropellin-3 | VWA domain containing CoxE-like protein (PF05762); von Willebrand factor type A domain (PF00092); von Willebrand factor type A domain 3 (PF13768); von Willebrand factor type A domain 2 (PF13519); EGF-like domain (PF00008); Human growth factor-like EGF (PF12661); Calcium-binding EGF domain (PF07645); Chitin binding Peritrophin-A domain (PF01607) | No | No | No |
| CGI_10000860 | 2.61154372 | 0.001486126479 | Unknown | - | No | No | No |
| CGI_10004609 | 2.59549451 | 0.000511252222 | Growth/differentiation factor 9 | Transforming growth factor beta like domain (PF00019) | No | No | No |
| CGI_10022464 | 2.19889547 | 0.003510355552 | Sushi, nidogen and EGF-like domain-containing protein 1 | Nidogen-like (PF06119); AMOP domain (PF03782); Sushi repeat (SCR repeat) (PF00084) | Yes (Zhang et al. 2012) | No | No |
| CGI_10026040 | -9.55560280 | 4.76515514e-05 | Unknown | - | No | No | No |
| CGI_10005508 | -7.26059669 | 5.40930022e-37 | No hit but it has PFAM domain | Dienelactone hydrolase family (PF01738) | No | No | No |
| CGI_10002171 | -5.21758417 | 4.68156100e-11 | Unknown | - | No | No | No |
| CGI_10013066 | -4.75444684 | 0.000439028973 | Complement C1q tumor necrosis factor-related protein 3 | C1q domain (PF00386) | No | Yes | No |
| CGI_10010209 | -4.63348063 | 0.000105936504 | Unknown | - | No | No | No |
| CGI_10024880 | -3.46101920 | 0.040670422330 | No hit but it has PFAM domain | WAP-type (Whey Acidic Protein) 'four-disulfide core' (PF00095) | No | No | No |
| CGI_10021348 | -3.44502005 | 0.003907284211 | cAMP-regulated D2 protein | Carboxylesterase family (PF00135); BD-FAE (PF20434); Abhydrolase_3 alpha/beta hydrolase fold (PF07859) | No | No | No |
| CGI_10002142 | -3.00929245 | 0.002101908985 | Unknown | - | No | No | No |
| CGI_10011370 | -2.58020742 | 0.001892727432 | Multiple epidermal growth factor-like domains protein 10 | - | No | No | No |
| CGI_10018833 | -2.20007387 | 0.012740818532 | Extracellular superoxide dismutase [Cu-Zn] | Copper/zinc superoxide dismutase (SODC) (PF00080) | Yes (Zhang et al. 2012) | No | No |

**Supplementary Table 3.** Gene Ontology terms (Level 2) of differentially expressed genes encoding secreted proteins in the mantle cells of oysters kept under normal (8.0 ± 0.2) and low (7.6 ± 0.2) pH conditions.

| **BIOLOGICAL PROCESS** | | | |
| --- | --- | --- | --- |
| **Gene Ontology Term** | **Gene Ontology Accession** | **Differentially expressed genes** | |
|  |  | **Up-regulated** | **Down-regulated** |
| Biological adhesion | GO:0022610 | CGI_10012073; CGI_10022464 | CGI_10011370 |
| Positive regulation of biological process | GO:0048518 | CGI_10015627; CGI_10025581; CGI_1000679 | CGI_10013066; CGI_10011370 |
| Biological regulation | GO:0065007 | CGI_10025581; CGI_10006796; CGI_10015627 | CGI_10013066; CGI_10018833 |
| Regulation of biological process | GO:0050789 | CGI_10006796; CGI_10025581; CGI_10015627 | CGI_10013066; CGI_10011370 |
| Response to stimulus | GO:0050896 | CGI_10015627; CGI_10025581; CGI_10012073; CGI_10023899; CGI_10002386 | CGI_10013066; CGI_10018833 |
| Cellular processes | GO:0009987 | CGI_10025581; CGI_10006796; CGI_10015627; CGI_10012073 | CGI_10013066; CGI_10011370; CGI_10018833 |
| Signaling | GO:0023052 | CGI_10006796; CGI_10025581; CGI_10015627 | CGI_10013066 |
| Negative regulation of biological processes | GO:0048519 | CGI_10006796; CGI_10025581 | CGI_10013066 |
| Metabolic process | GO:0008152 | CGI_10025581; CGI_10023899 | CGI_10013066; CGI_10018833 |
| Developmental process | GO:0032502 | CGI_10006796; CGI_10025581 | CGI_10011370 |
| Localization | GO:0051179 | CGI_10006796; CGI_10025581; CGI_10015627; CGI_10023899 | CGI_10013066; CGI_10011370 |
| Multicellular organismal process | GO:0032501 | CGI_10006796; CGI_10025581; CGI_10012073 | CGI_10013066; CGI_10018833; CGI_10011370 |
| Multi-organism process | GO:0051704 | CGI_10006796; CGI_10015627; CGI_10012073 |  |
| Locomotion | GO:0040011 | CGI_10006796 | CGI_10011370 |
| Detoxification | GO:0098754 |  | CGI_10018833 |
| Immune system process | GO:0002376 | CGI_10015627 |  |
| Cell proliferation | GO:0008283 |  | CGI_10011370 |
| Cellular component organization or biogenesis | GO:0071840 | CGI_10015627; CGI_10012073 | CGI_10011370 |
| Reproductive process | GO:0022414 | CGI_10006796 |  |
| Reproduction | GO:0000003 | CGI_10006796 |  |
| **MOLECULAR FUNCTION** | | | |
| **Gene Ontology Term** | **Gene Ontology Accession** | **Differentially expressed genes** | |
|  |  | **Up-regulated** | **Down-regulated** |
| Binding | GO:0005488 | CGI_10027000; CGI_10002386; CGI_10015627; CGI_10006796; CGI_10012073; CGI_10022464; CGI_10025581; CGI_10004609 | CGI_10013066; CGI_10011370; CGI_10018833 |
| Molecular function regulator | GO:0098772 | CGI_10004609; CGI_10006796 | CGI_10024880 |
| Molecular transducer activity | GO:0060089 | CGI_10015627 |  |
| Cargo receptor activity | GO:0038024 | CGI_10015627 | CGI_10011370 |
| Antioxidant activity | GO:0016209 | CGI_10002386 | CGI_10018833 |
| Catalytic activity | GO:0003824 | CGI_10002386 | CGI_10018833; CGI_10005508; CGI_10021348 |
| Structural molecule activity | GO:0005198 | CGI_10006796; CGI_10012073 |  |
| Transporter activity | GO:0005215 | CGI_10023899 |  |
| Toxin activity | GO:0090729 | CGI_10028906 |  |

**Supplementary Table 4.** Gene Ontology terms (Level 3) of differentially expressed genes encoding secreted proteins in the mantle cells of oysters kept under normal (8.0 ± 0.2) and low (7.6 ± 0.2) pH conditions.

| **BIOLOGICAL PROCESS** | | | |
| --- | --- | --- | --- |
| **Gene Ontology Term** | **Gene Ontology Accession** | **Differentially expressed genes** | |
|  |  | **Up-regulated** | **Down-regulated** |
| Cellular response to stimulus | GO:0051716 | CGI_10015627; CGI_10006796; CGI_10025581 | CGI_10013066; CGI_10018833 |
| Regulation of response to stimulus | GO:0048583 | CGI_10015627; CGI_10006796; CGI_10025581 | CGI_10013066 |
| Negative regulation of response to stimulus | GO:0048585 | CGI_10006796; CGI_10025581 | CGI_10013066 |
| Response to chemical | GO:0042221 | CGI_10006796; CGI_10015627 | CGI_10018833 |
| Response to stress | GO:0006950 | CGI_10002386; CGI_10023899; CGI_10015627; CGI_10025581 | CGI_10018833; CGI_10013066 |
| Response to abiotic stimulus | GO:0009628 | CGI_10023899 | CGI_10018833 |
| Response to external stimulus | GO:0009605 | CGI_10015627; CGI_10012073; CGI_10025581 | CGI_10013066 |
| Response to biotic stimulus | GO:0009607 | CGI_10015627; CGI_10012073 |  |
| Immune response | GO:0006955 | CGI_10015627 |  |
| Positive regulation of response to stimulus | GO:0048584 | CGI_10015627 | CGI_10013066 |
| Response to endogenous stimulus | GO:0009719 | CGI_10006796 |  |
| **MOLECULAR FUNCTION** | | | |
| **Gene Ontology Term** | **Gene Ontology Accession** | **Differentially expressed genes** | |
|  |  | **Up-regulated** | **Down-regulated** |
| Drug binding | GO:0008144 | CGI_10027000 |  |
| Carbohydrate derivative binding | GO:0097367 | CGI_10027000 |  |
| Protein binding | GO:0005515 | CGI_10025581; CGI_10004609; CGI_10006796; CGI_10022464 | CGI_10013066; CGI_10011370 |
| Ion binding | GO:0043167 | CGI_10015627; CGI_10006796; CGI_10012073; CGI_10022464; CGI_10027000 | CGI_10018833 |
| Amide binding | GO:0033218 | CGI_10027000 |  |
| Heterocyclic compound binding | GO:1901363 | CGI_10002386; CGI_10027000 |  |
| Organic cyclic compound binding | GO:0097159 | CGI_10002386; CGI_10027000 |  |
| Cofactor binding | GO:0048037 | CGI_10027000; CGI_10002386 |  |
| Sulfur compound binding | GO:1901681 | CGI_10027000 |  |
| Small molecule binding | GO:0036094 | CGI_10027000; CGI_10015627 |  |
| Carbohydrate binding | GO:0030246 | CGI_10015627 |  |
| Protein-containing complex binding | GO:0044877 | CGI_10015627 |  |
